# The presence of a G-quadruplex prone sequence upstream of a minimal promoter increases transcriptional activity in the yeast *S. cerevisiae*

**DOI:** 10.1101/2023.06.23.546269

**Authors:** Libuše Kratochvilová, Matúš Vojsovič, Natália Valková, Lucie Šislerová, Zeinab El Rashed, Alberto Inga, Paola Monti, Václav Brázda

## Abstract

Non-canonical secondary structures in DNA are increasingly being revealed as critical players in DNA metabolism, including modulating the accessibility and activity of promoters. These structures comprise the so-called G-quadruplexes (G4s) that are formed from sequences rich in guanine bases. Using a well-defined transcriptional reporter system, we sought to systematically investigate the impact of the presence of G4 structures on transcription in yeast *S. cerevisiae*. To this aim, different G4 prone sequences were modeled to vary the chance of intramolecular G4 formation, analyzed *in vitro* by Thioflavin T binding test and circular dichroism and then placed at the yeast *ADE2* locus on chromosome XV, downstream and adjacent to a P53 response element (RE) and upstream from a minimal *CYC1* promoter and Luciferase 1 (*LUC1*) reporter gene in isogenic strains. While the minimal *CYC1* promoter provides for basal reporter activity, the P53 RE enables *LUC1* transactivation under the control of the human P53 family proteins expressed under the inducible *GAL1* promoter. Thus, the impact of the different G4 prone sequences on both basal and P53 family proteins dependent expression was measured after shifting the yeast cells onto galactose containing medium. The results showed that the presence of G4 prone sequences upstream of a yeast minimal promoter can increase its basal activity proportionally to their potential to form intramolecular G4 structures; consequently, this improved accessibility, when present near the target binding site of P53 family transcription factors can be exploited in order to regulate the transcriptional activity of P53, P63 and P73 proteins.

## Introduction

Although the shape of the DNA molecule is primarily epitomized in the double helix, the ENCODE project has shown that DNA can form noncanonical secondary structures with a relevant biological significance ^1^. These structures include the so-called G-quadruplexes (G4s) that are formed from sequences rich in guanine bases. G4s are constituted of four guanine bases arranged in a square planar conformation (G-tetrad) held together by Hoogsteen hydrogen bonding and further stabilized by potassium or sodium ions ^2,3^. G4s can be presented in various topologies including antiparallel, parallel, and hybrid structures, depending on the relative orientation of the DNA strand within the structure; intermolecular G4 structures can also be formed when more than one DNA strand create the final structure ^4^. G4s have been shown to be involved in processes such as DNA gene replication, transcription, translation, and maintenance of genome stability; they occur in specific sequences with functional significance such as telomeres and promoter regions of oncogenes, and also on 5’ and 3’ untranslated regions (UTR) of messenger RNAs ^2,5,6^. Interestingly, bioinformatics analysis of the binding sites of several transcription factors showed their potential to interact with G4s ^6,7^ and provided the basis for the assumption that positive or negative regulations occur between transcription factors and G4 structures ^8^. Moreover, the increased frequency of G4 motifs in gene promoters compared to the rest of the genome suggests a significant contribution to the regulation of the expression of those genes ^9^. Experiments carried out with a single chain antibody specific for G4 structures showed the involvement of G4s not only during the initiation of transcription, but also during its termination ^8^. The first verified evidence of G4s influence on gene expression was demonstrated for the *MYC* oncogene, where mutations of a G4 in the promoter region affected *MYC* expression *in vivo* ^10,11^. Following studies addressed the regulation of transcription through G4 ligands (e.g., TMPyP4 and many others), resulting in a comparable decrease in transcription of *MYC* ^11^, *KRAS* ^12^ and *KIT* ^13^ oncogenes. Therefore, G4s targeting is suggested for cancer therapy ^14,15^.

The P53 family of transcription factors comprises the structurally related P53, P63, and P73 proteins that share an N-terminal transactivation domain (TA), a central sequence-specific DNA binding domain (DBD) and an oligomerization domain (OD) at C-terminus. All three proteins mainly act in the cells as tetramers able to induce the expression of a plethora of target genes involved in different cellular pathways, including cell proliferation, apoptosis, DNA repair, angiogenesis, metabolism and differentiation ^16–18^. The three proteins are also characterized by a similar gene structure that produces groups of mRNAs controlled by separate promoters and encoding proteins with alternative N-terminal regions ^19,20^. While the variants generated from the external promoter contain the complete TA domain (TA-isoforms) and are transactivation competent, those generated from internal promoter lack the full TA domain (ΔN-isoforms) but still bind DNA, showing both specific transactivation ability and dominant negative activity. To further complicate the P53 family landscape, the transcripts are also subject to alternative splicing of the C-terminal portion, giving rise to a combinatorial variety of P53, P63 and P73 specific isoforms ^21,22^.

Transcriptional regulation by P53 family proteins is mainly achieved through the binding to a degenerate DNA motif known as P53 response element (RE) consisting in two decameric half-sites separated by a short base spacer (RRRCWWGYYY-n-RRRCWWGYYY, where R stands for a purine, W for A/T, Y for a pyrimidine and n for spacer) ^23,24^. However, while binding affinities of the P53 family transcription factors are related to primary sequence features of the RE, other factors modulate the binding and the transactivation that can be related both to DNA accessibility within chromatin and to various features of the so-called indirect-readouts or shape-readouts; these are dependent on DNA structural features that can be influenced by trans-factors including proteins, non-coding RNAs, epigenetic changes, and can influence nucleosome density and positioning ^24^.

Interestingly, local unfolding events of the DNA double helix are able to stimulate the formation of G4s structures, potentially contributing to orchestrate P53 family cell-type specific transcriptomes ^25^. Based on the critical role of these DNA structural elements in modulation of transcriptional activity, we previously evaluated the influence of a G4 prone sequence from KSHV (Kaposi sarcoma-associated herpes virus) in the proximity of a P53 RE from *BBC3* (PUMA) target gene on the transactivation potential of ΔN- and TA-variants of P53 family α isoforms ^25,26^. Here, by using the same yeast-based assay, we study the effect of different G4 prone sequences both on basal and P53 family dependent expression of the Luciferase 1 reporter gene (*LUC1*). To this aim, different G4 prone sequences were modeled to vary the chance of intramolecular G4 formation and studied *in vitro* for the different propensity of forming G4 structures by Thioflavin T binding and circular dichroism analyses. The sequences were then placed downstream and adjacent from the PUMA RE, constructing otherwise isogenic yeast reporter strains. The impact of various G4 prone sequences was measured after transforming the cells with centromeric inducible expression vectors and shifting them to galactose-containing medium to modulate the expression levels of the P53, P63, and P73 transcription factors (wild-type TA α isoforms). The results obtained highlight that the higher is the propensity to form G4 structures upstream of the promoter, the higher is the transcriptional activity both at basal and transcription factor-dependent level.

## Material and Methods

### Synthetic oligonucleotides

Synthetic oligonucleotides were purchased from Sigma-Aldrich and diluted with ultrapure water to the final concentration of 100 μM (**Table 1**). The PUMA sequence was derived from the *BBC3* (PUMA) P53 target gene; oligonucleotide sequences with different G4s formation potential were designed to decrease the chance of intramolecular G4 formation by mutating the KSHV G-quadruplex sequence ^25^. KSHV-Mut2.0 and KSHV-Mut1.5 sequences were designed by the G4 Killer program ^27^ to change G4 Hunter score below 2.0 and 1.5, respectively; the sequences of KSHV-1NO, KSHV-2NO and KSHV-3NO were designed by sequential substitution of G bases in guanine repeats. The propensity of the G4 formation of used sequences were predicted by the G4 Hunter program and measured as G4 Hunter scores (**Table 1**) ^28^.

**Table 1.**
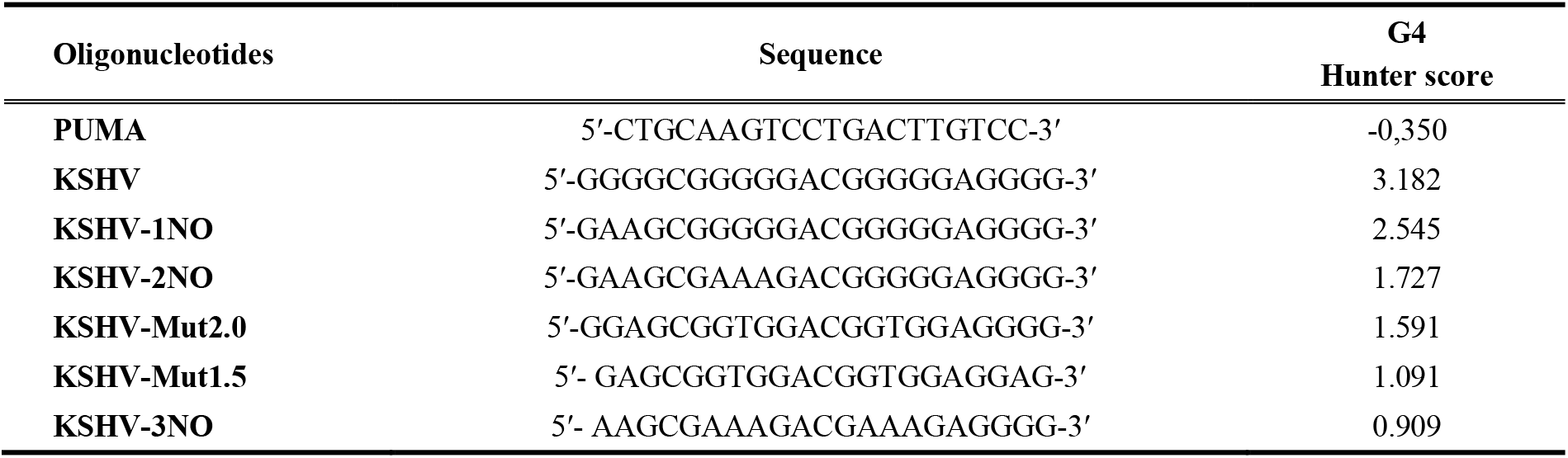
Oligonucleotides used in present study.

### Thioflavin T assay analysis

The thioflavin T (ThT) binding test was performed according to the procedure described in ^29^. Synthetic oligonucleotides were denatured in 100 mM Tris-HCl (pH 7.5) or 100 mM Tris-HCl (pH 7.5) containing 100 mM KCl at a final concentration of 2 μM (95°C for 5 minutes) and gradually cooled to laboratory temperature. Oligonucleotides were mixed with ThT in 2:1 molar ratio and measured in a 384-well titration microplate reader at room temperature in three replicates (excitation at 425 nm). Fluorescence emission was detected at 490 nm; the measured fluorescence intensities of individual oligonucleotides were related to the fluorescence intensity of the buffer with ThT without DNA (I/I_0_).

### Circular Dichroism spectroscopy analysis

Synthetic oligonucleotides were denatured in 10 mM Tris-HCl or 10 mM Tris-HCl containing 100 mM KCl at final concentration 3 μM (95°C for 5 minutes) and gradually cooled to room temperature. Circular Dichroism (CD) allows the characterization of the topology of the resulting secondary structure by comparison with model spectra ^30,31^. Spectra were measured on a Jasco 815 dichrograph in 1 cm tapered quartz cuvettes that were placed in a thermostatically controlled holder at 20°C. Four scans of each sample were taken at a scan rate of 100 nm·min^-1^ with a data pitch of 0.5 nm in the wavelength range 200–330 nm, averaged, and the resulting spectra were smoothed using the Savitzky-Golay smoothing algorithm with a 15-point window. The CD signal was expressed as the difference in the molar absorption coefficient Δε of the left- and right-hand polarized light. The CD measurements were taken also after 24 hours.

### Yeast reporter strains, plasmids and manipulations

*S. cerevisiae* reporter strains that contain, in addition to the P53 RE from the *BBC3* (PUMA) target gene (5′-CTGCAAGTCCTGACTTGTCC-3′), the different G4 prone sequences (**Table 1**), were already available ^25^ or newly created by the *Delitto Perfetto* approach ^32^. G4 prone sequences were located downstream of the RE and immediately up-stream of the minimal CYC1-derived promoter controlling the expression of the *LUC1* reporter gene, cloned at the yeast *ADE2* locus on chromosome XV ^33^. The following isogenic yeast strains yLFM-PUMA (PUMA), yLFM-PUMA-KSHV (P-K), yLFM-PUMA-KSHV-1NO (P-K-1NO), yLFM-PUMA-KSHV-2NO (P-K-2NO), yLFM-PUMA-KSHV-Mut2.0 (P-K-Mut2.0), yLFM-PUMA-KSHV-Mut1.5 (P-K-Mut1.5) and yLFM-PUMA-KSHV-3NO (P-K-3NO) strains were used. Vectors based on pTSG plasmid containing the coding sequences for wild-type P53 family TA α isoforms (i.e., P53, P63, and P73) expressed under a yeast-inducible *GAL1,10* promoter were already available along with the empty vector pRS314 (*TRP1* selection marker) ^34^. Yeast cells were transformed by the lithium acetate method as previously described ^35^. Yeast transformants were selected on plates lacking tryptophan and containing high amounts of adenine (200mg/L), given the deletion of *ade2* of all strains, and incubated for 3 days at 30°C. Colonies were then patched onto the same selective plates to expand them prior to the reporter assay.

### Yeast luciferase assay

Yeast transformants (empty and wild-type P53 family TA α isoforms) were resuspended using a 96-well plate in synthetic, tryptophan-selective liquid medium containing raffinose as carbon source and then diluted in the selective media supplemented with the amount of galactose needed to reach the selected final concentrations (0.016% and 1%). The shift from glucose containing plates to raffinose plus galactose media enables the activation of the *GAL1,10* promoter and the induction of the expression of P53 family proteins at moderate and high levels ^33^. The reporter expression took place after 6 hours of incubation time at 30°C. Luciferase assays were performed using a white 96-well plate, where 20 μl of yeast culture was mixed in each well with an equal volume of 2x Passive Lysis Buffer (Promega) to permeabilize cells by 15 minutes of shaking at room temperature; then the firefly luciferase substrate (20 μl) was added (Bright-Glo™ Luciferase Assay Kit, Promega). Light unit values were normalized to the OD_600_ absorbance of each culture that was measured from the 96-well culture plates. Results were expressed by: i) relative light units (RLU, i.e., Firefly units/OD), ii) fold changes using as reference the RLU values obtained from transformants with the empty pRS314 vector and iii) relative activities as ratio of fold changes. For each experiment, at least three independent cultures were measured.

### Statistical analysis

Ordinary One-way anova followed by comparisons test were performed by using GraphPad Prism version 9 for Mac, GraphPad Software, San Diego, California, USA.

## Results

### Formation of G4s structures *in vitro*

Firstly, we assessed G4s formation in oligonucleotides (**Table 1**) using the ThT assay; as the G4 structure is stabilized by potassium ions, we used buffers with and without KCl addition. In both buffers, the samples with the potential for the formation of G4 achieved a fluorescence intensity several times higher than the PUMA oligonucleotide (negative control), whose fluorescence reached the value of the ThT background signal value (**Figure 1A, B; Supplementary Table 1**); furthermore, the fluorescent signals were higher in the buffer with potassium ions (**Figure 1A, B; Supplementary Table 1**).

**Figure 1.**
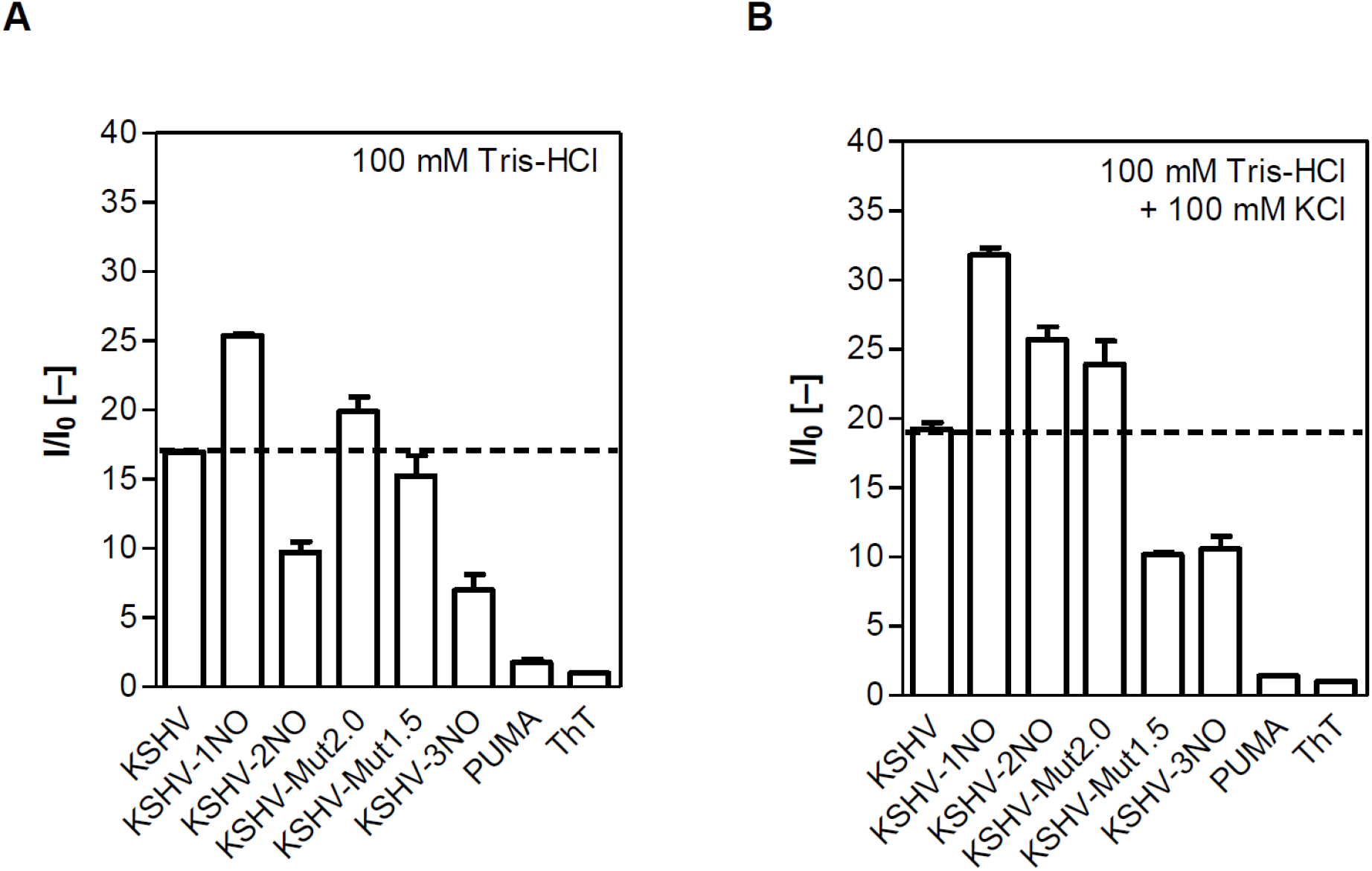
Evaluation of G4s formation potential in vitro by ThT assay. Oligonucleotides were hybridized in 100 mM HCl **(A)** or in the same buffer with the addition of 100 mM KCl **(B)**. The I_(KCl)/_I_(0)_ fold values are reported in Supplementary Table S1.

The fluorescence for KSHV-1NO, KSHV-2NO and KSHV-3NO oligonucleotides decreased in accordance to the decrease of the G4 Hunter score (**Figure 1A, B; Supplementary Table 1**); the increase in the fluorescent signal in presence of potassium ions was most pronounced on the KSHV-2NO oligonucleotide, while the lowest increase was observed by the KSHV-1NO sequence. The KSHV-3NO oligonucleotide, characterized by the lowest G4 Hunter score, reached approximately 1.5 times higher fluorescence intensity than the one hybridized without KCl; this increase by a G4 sequence that is designed to strongly reduce the formation of the intramolecular G4 structure is probably associated with the ability of potassium ions to facilitate the formation of intermolecular G-quadruplex structures.

KSHV-Mut2.0 and KSHV-Mut1.5 oligonucleotides showed comparable fluorescence intensity in buffer without potassium ions (**Figure 1A; Supplementary Table S1**). A higher fluorescence intensity in KCl-containing medium was achieved only by the KSHV-Mut2.0 sequence, which confirmed the formation and stabilization of G4 structures; conversely, the KSHV-Mut1.5 sequence showed a decrease in signal, with a value comparable to KSHV-3NO (**Figure 1B; Supplementary Table 1**). This result may suggest a destabilization of the emerging G4 structures or the formation of different loops through the contribution of electrostatic interactions with a less stable conformation.

Then, we characterized the G-quadruplex formation using CD spectroscopy analysis; measurements were also taken after 24 hours to determine changes in conformation over time (Figure 2; Figure S1).

**Figure 2.**
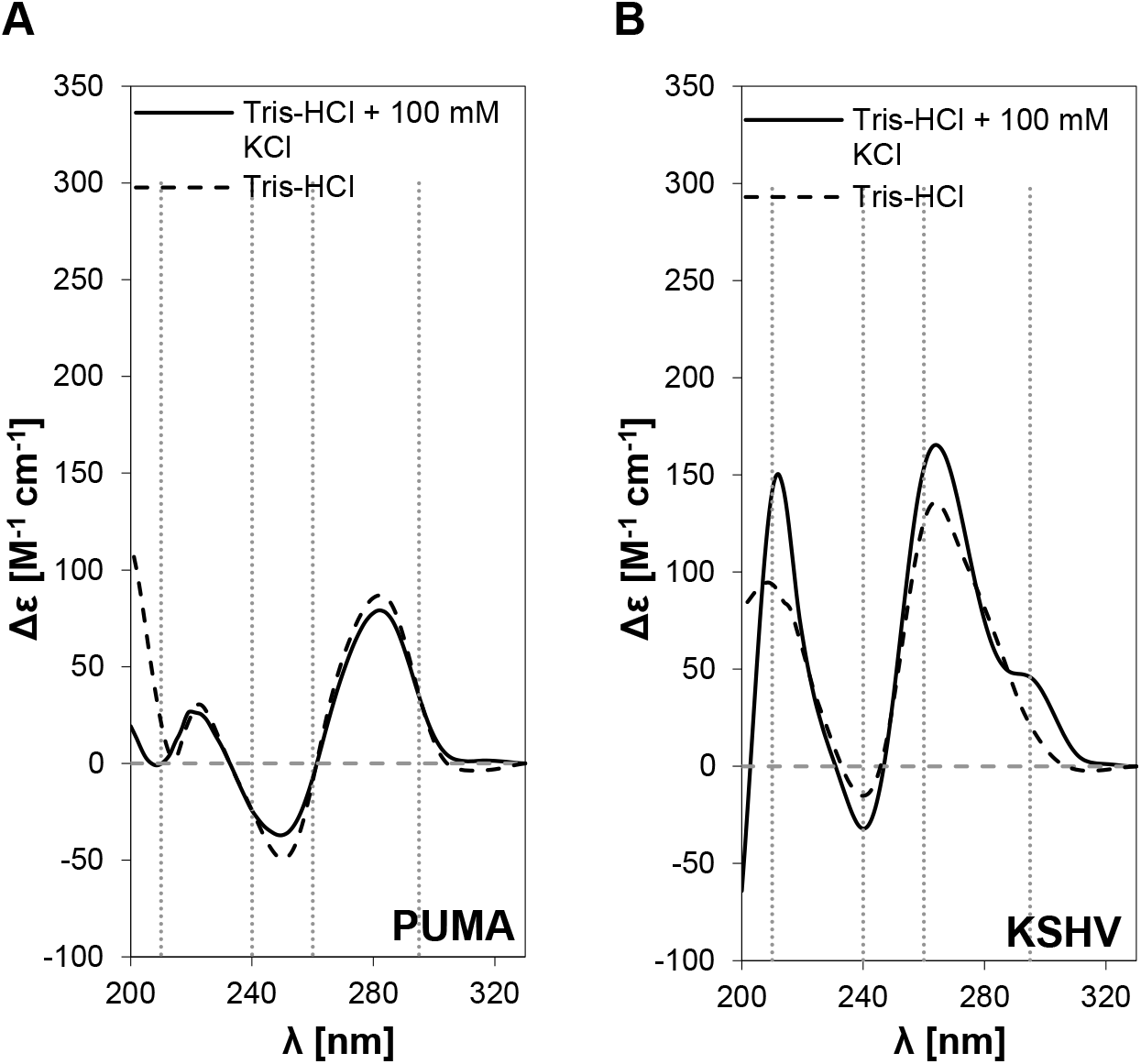
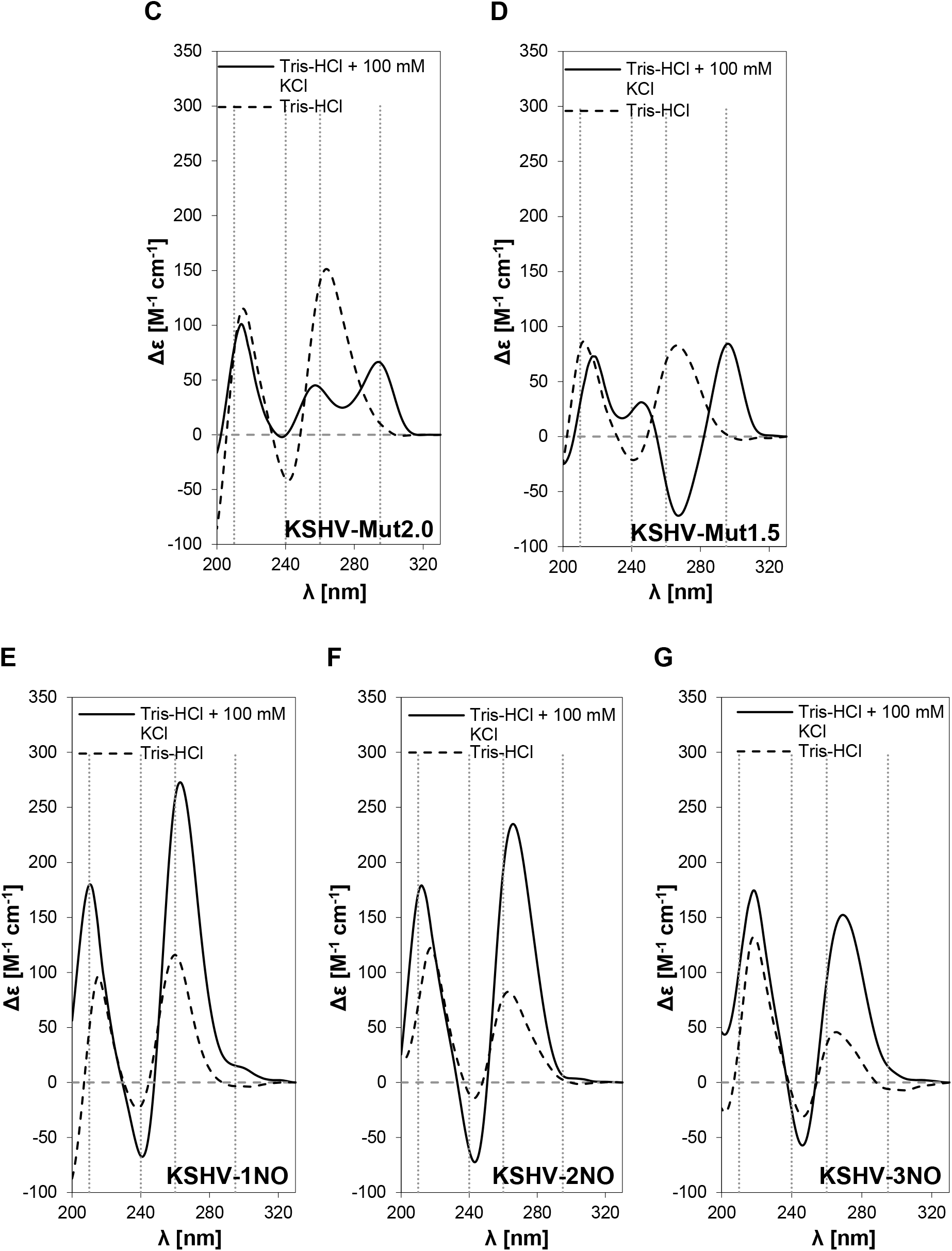
Evaluation of G4s formation potential *in vitro* by CD spectra analysis. CD spectra of oligonucleotides of **(A)** PUMA, **(B)** KSHV, **(C)** KSHV-Mut2.0, **(D)** KSHV-Mut1.5, **(E)** KSHV-1NO, **(F)** KSHV-2NO and **(G)** KSHV-3NO in medium without stabilizing potassium ions (dashed line) and in medium supplemented with 100 mM KCl (solid line). Wavelengths (210, 240, 260 and 295 nm) characteristic for the presence of G4 conformations in CD spectra are highlighted by gray dashed lines.

PUMA oligonucleotide spectra (**Figure 2A)** did not show any typical peaks for G4 even after the addition of potassium ions, as apparent from the absence of the G4 characteristic peak in the 210 nm region ^36^; moreover, the spectra were nearly identical in both types of buffers, reaching positive peaks at 220 and 280 nm and a negative peak at 250 nm. Similarly, KSHV-3NO sequence with the lowest G4 Hunter score (0.909) did not form a G4 structure even in the presence of potassium ions (**Figure 2G)**. In contrast to PUMA and KSHV-3NO oligonucleotides, the KSHV sequence showed characteristics typical for a parallel G-quadruplex with the positive peaks around 210 and 260 nm and a negative peak at 240 nm; moreover, a small peak around 295 nm suggested that mixed or antiparallel G4 structures were also possible for this sequence (**Figure 2B)**. The strong formation of G4 structures in the KSHV sequence was also supported by the presence of the typical peaks described above even without potassium ions.

The spectra of the KSHV-1NO (**Figure 2E)** and KSHV-2NO (**Figure 2F)** suggested that these sequences could form a parallel G4 structure, especially in buffer containing potassium ions. The CD spectra of the KSHV-Mut2.0 (**Figure 2C)** and KSHV-Mut1.5 sequences (**Figure 2D)** reached positive peaks around 216 and 264 nm and a negative peak at 242 nm, indicating the possible presence of a parallel G4 structure without the addition of potassium ions; in potassium buffer, the latter oligonucleotide formed a hybrid G4 structure, characterized by spectra with positive peaks around 210, 260 and 295 nm.

After 24 hours for oligonucleotides KSHV-1NO, KSHV-2NO, KSHV-Mut2.0, and KSHV-Mut1.5 hybridized in KCl buffer, there was a slight increase in signal intensity without any change in topology; for the remaining motifs, there was a decrease in signal, and in some cases deviations in the measured spectra, but no evidence of a change in conformation over time (**Figure S1**). All together the results confirmed the differential propensity of oligonucleotides we selected to form G4 structures *in vitro*, revealing a certain consistence with the results of their selection based on the G4 Hunter score.

### Effect of the presence of G4 prone sequences on basal reporter activity in a yeast-based assay

The same panel of G4 prone sequences assayed *in vitro* were placed downstream and adjacent to PUMA RE in isogenic yeast reporter strains and analyzed by a functional assay. We began by examining the impact of the sequences on the basal expression of the *LUC1* reporter six hours after shifting the yeast cells to 0.016% and 1% galactose containing medium.

By plotting the RLU data and ordering the G4 prone sequences by the decreasing G4 Hunter score, a proportional trend was observed. G4 prone sequences KSHV and KSHV-1NO led to higher transcription and this effect was progressively reduced by decreasing the strength of the G4 sequence measured by the G4 Hunter score (**Figure 3**), being partially affected also by the type of G4 prone motif. Indeed, transcription in P-K-1NO, P-K-2NO and P-K-3NO strains decreased; similarly, this effect was evident with P-K-Mut2.0 and P-K-Mut1.5 strains. However, the P-K-3NO strain, characterized by the lowest G4 Hunter score (0.909) did not show the lowest RLU value, being in any conditions statistically different from the control PUMA. Conversely, the transcription in P-K-Mut1.5 strain (G4 Hunter score 1.091) was not or weakly statistically different from the one in PUMA at 0.016% and 1% galactose, respectively; lastly, the transcription in P-K and P-K-1NO strains was comparable and statistically higher (p<0.0001) than the one in PUMA.

**Figure 3.**
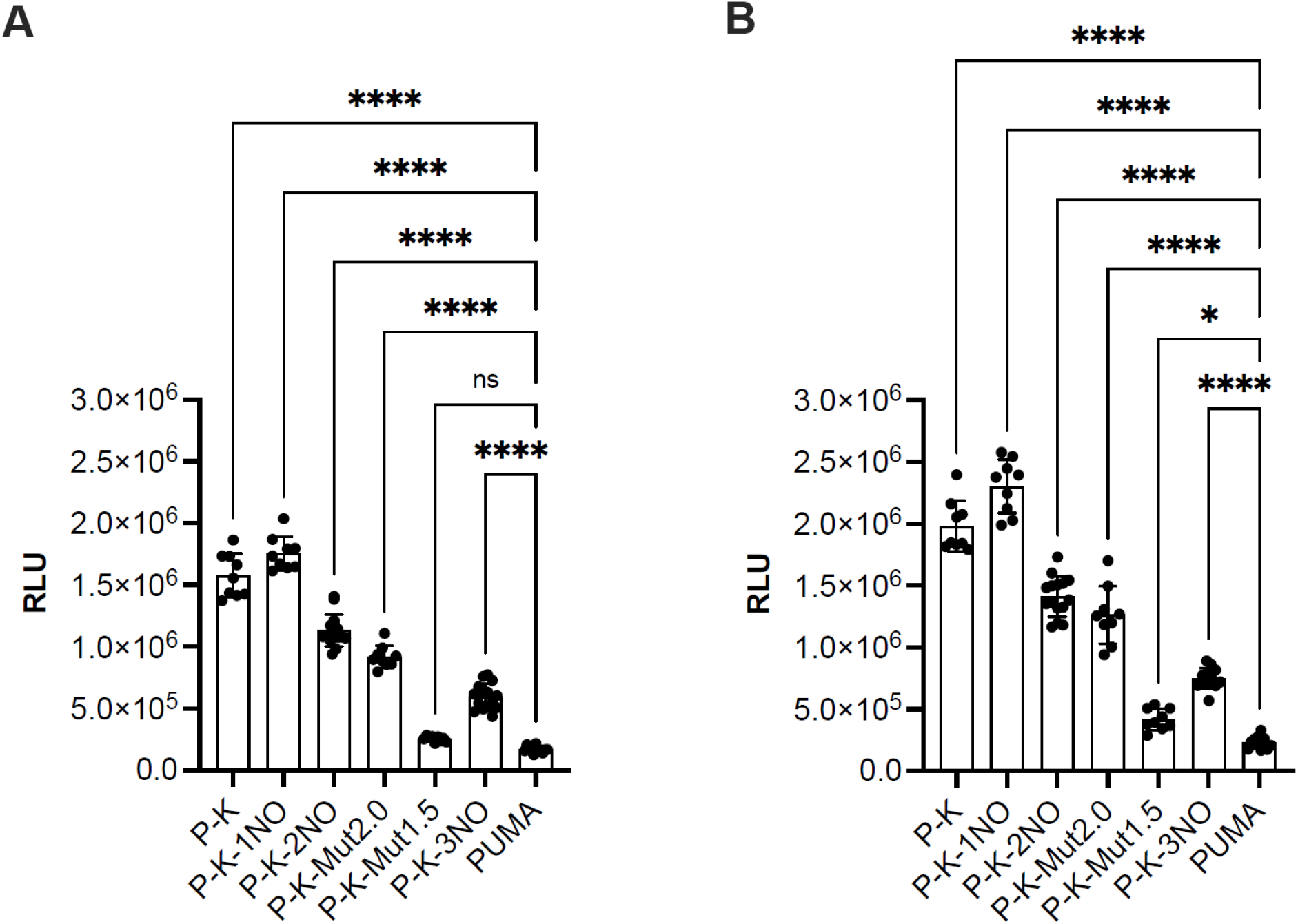
Effect of G4 prone sequences on basal reporter activity in the yeast *S. cerevisiae*. **(A)** RLU measurements for the indicated panel of yLFM reporter strains from empty plasmid (pRS314) yeast transformants at 0.016% Galactose for 6 hours. **(B)** RLU measurements as above at 1% Galactose for 6 hours. Data are presented as mean ± standard deviation, (SD) of at least three biological replicates, and individual values are also plotted. The symbols * and **** indicate significant differences with p=0.0461 and p<0.0001, respectively between PUMA strain and those containing G4 regulatory elements. ns, not significant. Ordinary one-way ANOVA test.

The results suggested that the basal reporter transcription can be stimulated by the presence of a G4 sequence in the yeast *S. cerevisiae* and in relationship to the propensity to form G4 structures.

### Effect of the presence of G4 prone sequences on transcription factor-dependent reporter activity

Next, we examined the potential impact of G4 prone sequences on transcription in a condition where sequence-specific transcription factors can bind upstream of the promoter and the G4 site, stimulating *LUC1* reporter expression. We exploited wild-type human P53 family proteins (*i*.*e*., TA P53, P63, and P73 α isoforms) under an inducible promoter to obtain moderate or high-level of their expression. As expected, the expression of the *LUC1* reporter was clearly enhanced already at the lower level of galactose inducer, especially by P53 protein with respect to basal transcription (**Figure 4**). The presence of G4 prone sequences with different scores had a weak effect on modulation of P53 transactivation (**Figure 4A, D)**; conversely, a trend, indicating that the P63- and P73-dependent reporter transcription was stimulated by the presence of G4 sequences, is present (**Figure 4B, E: P63; Figure 4 C, F: P73)**. Based on these results, we concluded that a strongly active transcription factor, such as wild-type P53 can be less influenced by the presence of the G4 prone sequences compared to the weaker P63 and P73 proteins.

**Figure 4.**
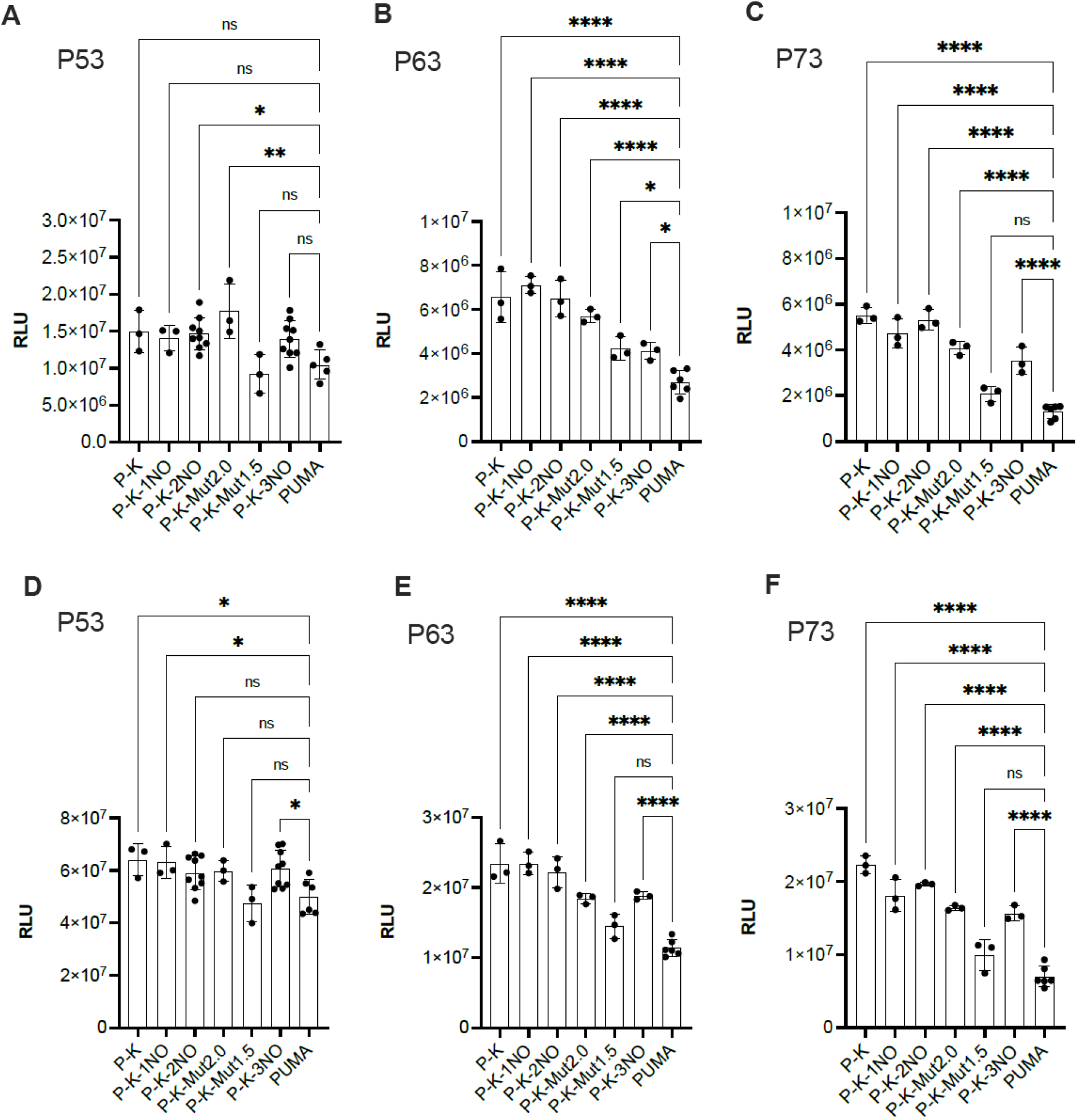
Effect of G4 prone sequences on P53 family-dependent reporter activity in the yeast *S. cerevisiae*. **(A-C)**: RLU measurements for the indicated panel of yLFM reporter strains expressing respectively P53, P63, and P73 at 0.016% Galactose for 6 hours. **(D-F)** RLU measurements as above at 1% Galactose for 6 hours. Data are presented as mean ± standard deviation (SD) of at least three biological replicates, and individual values are also plotted. The symbols *, ** and **** indicate significant differences with p ≤0.0406, p=0.0017and p<0.0001, respectively between PUMA strain and those containing G4 regulatory elements. ns, not significant. Ordinary one-way ANOVA test.

Then, by plotting the data as fold change, *i*.*e*., using the basal transactivation levels to normalize the impact of P53 family proteins expression in each strain, as previously done in our transactivation studies ^25,26^, lower fold change values for the G4-containing strains compared to the PUMA control were mostly evident (**Figure S2**). Overall, the results were consistent with the view that G4 prone sequences are associated with higher basal transcription, a feature that may favor the transactivation function of the weak transcription factors P63 and P73, which have reduced potential to interact with the basal transcription machinery.

This effect becomes more evident when focusing on the relative activity of P63 and P73 transcription factors with respect to P53 (**Figure 5)**. In general, the presence of G4 sequences with higher score reduces the gap in transactivation between P53 and the other two family members P63 and P73.

**Figure 5.**
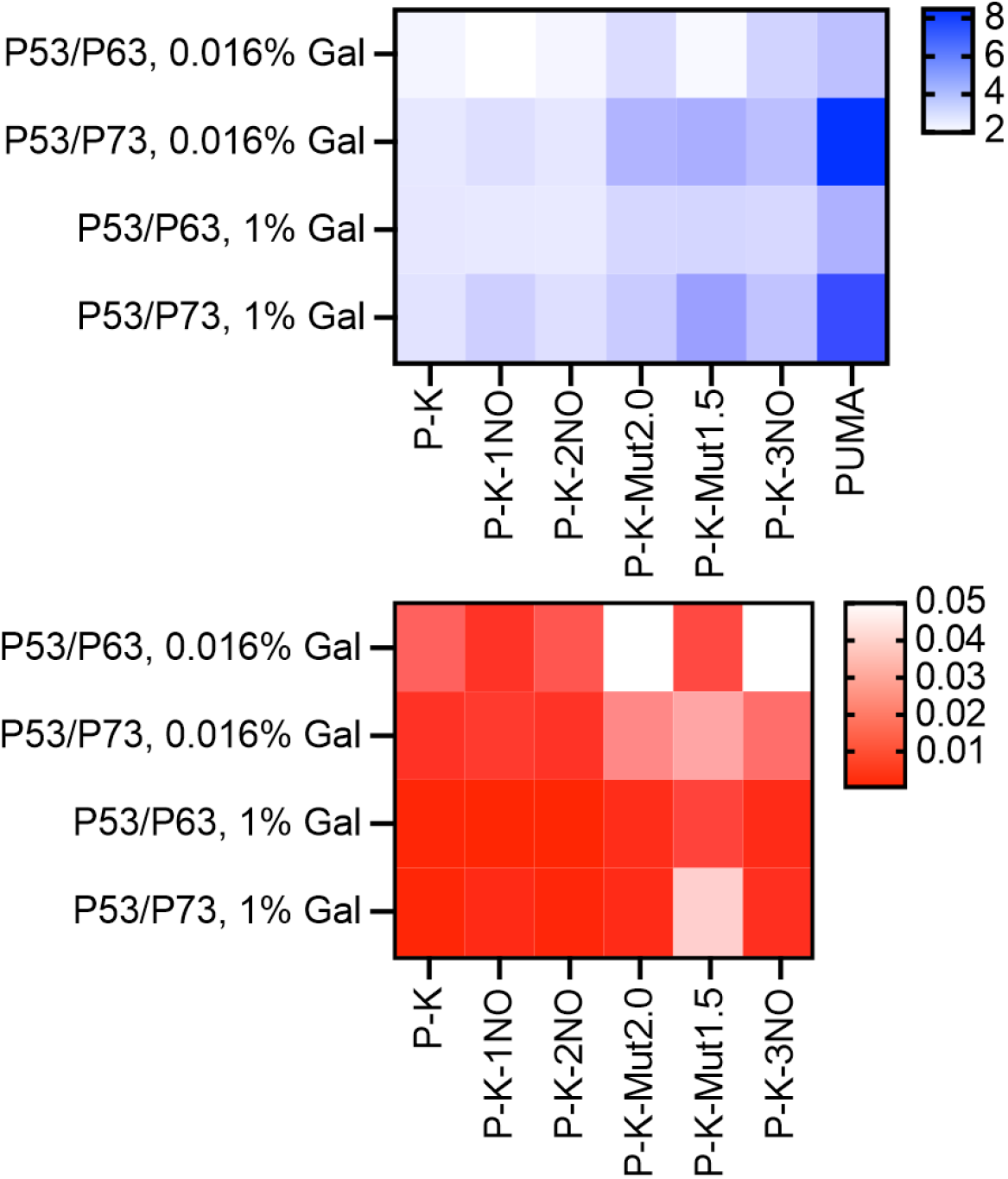
Effect of G4 forming sequences on P53/P63 and P53/P73 relative activity plotted as a heat map. The heatmap on the top presents the relative activity of P63 and P73 compared to P53 (shades of blue with the indicated color scale; average of three to six replicates) in the indicated panel of yLFM reporter strains; the heatmap on the bottom presents the results of multiple unpaired t-tests, assuming individual variance for each row and a two-stage step-up method to compare the relative activities measured with the control PUMA strain with those of the strains containing the other G4 regulatory elements (shades of red, p value <0.05; white = not significant.). Data with corresponding statistical analysis were obtained after 6 hours of growth in media containing galactose 0.016% or 1%. Data is also presented as bars graph in **Figure S3**.

## Discussion

Transcription is regulated by several mechanisms, one of the most important being proteins binding to promoters and enhancers ^37^. Sequence-specific transcription factors interacts with their target binding sites, but for many transcription factors it has been demonstrated that their specificity and activity can be influenced by the presence of non-B and higher order DNA structures ^38–41^. A prominent category of such structural elements includes DNA G-quadruplexes (G4s) consisting of secondary structures formed by stacked G-tetrads ^9,42^. G4 prone sequences are present in all organisms including *S. cerevisiae* ^43^ and human genome ^44–46^. Recently G4 structures were confirmed and visualized within cells using conformation antibodies or small molecule ligands, and their presence have been correlated with transcriptional regulation ^47,48^. Many studies support the view that G4s are favored by some transcription factors, affecting the transcriptional response ^49,50^. While several G4 binding proteins were evaluated in animals, including humans, the knowledge about G4-binding proteins in yeasts is limited and the influence of G4s on yeast transcription has not been deeply evaluated.

In this study, our objective was to develop a proof-of-concept framework to investigate how features of different G4 prone sequences can impact both basal and induced transcription in a defined, isogenic yeast model. While our approach relies on an artificial promoter-reporter construct, it has the advantage of testing defined variants of G4, embedded at the same position in a chromosomal locus upstream of a minimal promoter driving the expression of the quantitative *LUC1* reporter ^25,26^. Further upstream of the G4 motif is present a constant transcription factor RE from the *BBC3*(PUMA) target gene that can be targeted by the human P53 family proteins. Exploiting a tunable promoter, we were able to obtain results with a defined matrix of variables consisting of i) six different G4-forming proneness and a control sequence (i.e., PUMA) (**Table 1**), and ii) two different expression levels of the three transcription factors P63, P73, P53 (TA α variant) that differ for their binding affinity to the RE. The choice of P53 family transcription factors to investigate the impact of G4 prone sequences on induced transactivation was linked to our previous studies in which their transcriptional responses to many different variants of binding sites in the same chromosomal locus and reporter yeast system have been characterized ^33^. The possibility to test the entire family of P53 transcription factor was an added value of our proof-of-concept matrix; in fact, P53 family proteins can recognize binding sites with similar but not identical features. Also, the three proteins differ in the strength of transactivation that is in part related to differences in quaternary structures, and the presence of a different combination of domains ^22,51,52^.

Moreover, we chose the PUMA derived RE due to its intermediate transactivation potential, reasoning that it could represent a sensitive tool to monitor the impact of changes in the surrounding sequences ^25,33,53^. In fact, we have previously shown that high-affinity consensus P53 binding sites can lead to high-level transactivation even when P53 protein expression is kept at minimum levels, preventing the tuning of promoter responses ^54,55^. Indeed, the PUMA RE features three mismatches from the optimal P53 consensus site and does not contain the optimal, more flexible, CATG element in the so-called core CWWG motifs; at the same time, it is responsive also to P63 and P73 proteins.

The six G4 sequences selected in the present study were modeled on the positive control sequence derived from the KSHV virus sequence ^25^; specific changes were introduced to decrease the propensity to form G4s by the G4 Killer program ^27^ or by sequential substitution of G bases in guanine repeats. The resulting sequences were classified according to the G4 Hunter algorithm ^28^, a powerful and widely used tool for G4 prediction that takes into account G-richness and G-skewness of a DNA or RNA sequence and provides a quadruplex propensity score.

Firstly, the G4 prone sequences we selected were biophysically characterized by ThT assay and by CD spectroscopic analysis. ThT is a fluorescent probe that binds G4s much better compare to duplex or single stranded DNA and for which a procedure for the rapid detection of G4 structures has been developed and verified ^56^. By the ThT assay, we confirmed the formation of G4s not only in the previously analyzed KSHV sequence ^25^, but also in the oligonucleotides we selected and characterized by a lower G4 Hunter score (**Figure 1, Supplementary Table 1**). However, some deviations from the bioinformatic prediction of G4 formation were observed, mainly due to possibility of parallel G4 formation in short oligonucleotides. Indeed, the value of the G4 Hunter score is based on the number of concurrent G bases (G-runs) and the total length of the sequence. Although the presence of G-runs is essential for G4 formation, differences in bioinformatics-determined PQS (potential quadruplex-forming sequences) and preferences for G4 formation *in vitro* have been demonstrated ^57^. The results may have been influenced by the nature of ThT binding to DNA. A significant increase in fluorescence emission can occur due to the inhibition of the rotation of the ThT inner segments; binding of ThT to different DNA structures leads to different degrees of inhibition of torsional motion ^58^. Three different binding modes to DNA have been described for ThT: binding to the DNA cavities, intercalation between DNA bases, and external binding to the DNA phosphate groups; then, ThT bound in each mode has a different yield of fluorescence ^59^.

The results of CD spectroscopy analysis confirmed in the presence of potassium i) the stabilization of parallel G4 conformation in KSHV, KSHV-1NO, KSHV-2NO oligonucleotides, ii) an antiparallel conformation in KSHV-Mut1.5 sequence and iii) a hybrid conformation in KSHV-Mut2.0 motif. The fact that parallel G4 conformation is represented to a higher extent in KSHV-1NO and KSHV-2NO (**Fig. 2C, D**) sequence than in the positive control KSHV oligonucleotide **(Fig. 2B)**, where the positive peak at 295 nm in the CD spectra partially indicates the formation of hybrid G4, is consistent with the results of the ThT assay. In fact, the ThT molecule was reported to possess high specificity for the parallel G4 conformation ^60,61^. The change in the G4 conformation of KSHV-Mut1.5 also corresponds to the measured results from the ThT assay, when a decrease in fluorescence intensity was observed in the potassium ion environment, indicating the binding of ThT to the less preferred G4 structure.

Concurrently, transactivation data in our yeast-based experimental system indicate that the presence of these G4 prone sequence results in significant enhancement in basal transcription that is proportional to the potential to form secondary structures (**Figure 3**). Higher levels of transactivation are also apparent when the transcription rates are enhanced by the expression of the sequence-specific P53 family transcription factors, but in this case an inverse relation to affinity and transactivation potential is evident (**Figure 4**).

Transcription factor induced activity is typically expressed in reporter assays as fold change transactivation, to focus on the direct impact of a trans-factor on gene expression. Considering that the promoter-reporter system used reached a maximal rate of transactivation, the relative fold change in expression due to the action of the P53 family proteins is lower when a G4 prone sequence is embedded near the promoter site, due to the effect of these sequence in increasing the basal, P53-independent transcription. Fold change graphs show a negative impact of G4 prone sequences particularly for P53 (**Figure S2**); indeed this negative effect is proportional to the G4 Hunter score, with the partial exception of KSHV-3NO. However, we wish to emphasize that the fold change view is partially misleading since the presence of G4s actually increase overall promoter transcription in all cases, but particularly the basal activity. In general, P53 exhibits in yeast a much stronger transactivation potential than P73 and P63 as TA α variant ^54,62^. We exploited this feature, along with the inducible and tunable nature of the expression vector, to ask if the impact of studied G4 prone sequences could be more relevant in cooperating with the weaker transcription factors P63 and P73. This seemed indeed to be the case and led to a significant reduction in the relative activity of P53 over that of P63 and P73 with the presence of G4 forming sequence at the promoter site and inversely proportional to the G4 Hunter score (**Figure 5**).

While we have no formal proof that G4 sequences are stably formed at the endogenous yeast locus where they were cloned, the impact of those sequences on transactivation broadly followed the predictions from the G4 Hunter score in a direct manner. However, there were some exceptions, in fact the KSHV-1NO sequence showed an equivalent or even a slightly higher transactivation compared to the highest-scoring KHSV sequence; this is in part consistent with the results of ThT measurements performed in the presence of KCl. Moreover, the KSHV-3NO sequence showed higher basal transactivation than the higher scoring KSHV-Mut1.5 motif, although *in vitro* experiments did not show propensity for the former sequence to form G4s. One limitation from our *in-cellulo* approach, is represented by the inability to fully understand the contribution of a G4 prone sequence as primary sequence motifs or the effect at the level of local DNA structures. In other words, we cannot exclude that, by replacing the G content with A stretches in the oligonucleotides, new binding sites for yeast resident transcription factors are formed that can lead to higher basal transactivation independently from the formation of a G4 structure.

In conclusion, our proof-of-concept experiment modeled the impact of defined changes in G4 prone sequences at a yeast chromatin locus on both basal and induced transactivation, exploiting an isogenic well-established reporter assay. Results support the view that G4s cooperate with transcription factors and the basal transcription machinery. The cooperation is more relevant for the relatively weaker transcription factors P63 and P73 than for the highly responsive P53 protein. The fact that locally present G4 structures can change the functionality of P53 family proteins, could have important implications for the functional classification of pathogenetic P53 or P63 missense mutant alleles that retain partial transactivation capacity, therefore influencing their relative activity.

## Supplementary Materials

Table S1. Fluorescence intensity I/I_0_ determined from oligonucleotides with G4 formation potential.

Figure S1. CD spectra of oligonucleotides under study at T_0_ (left panel) and T_24_ (right panel)

Figure S2. Effect of G4 prone sequences on P53 family transactivation.

Figure S3. Effect of G4 forming sequences on P53/P63 and P53/P73 relative activity plotted as bar graphs.

## Author contributions

The authors confirm contribution to the paper as follows: study conception and design (VB, PM, AI); experimental data collection (LK, NV, MV, LŠ, ZE, PM), analysis and interpretation of the results (LK, MV, NV, LŠ, ZE, AI, PM, VB); draft manuscript preparation (LK, MV, AI, PM, VB). All authors reviewed and approved the final version of the manuscript.

## Funding

This work was supported by the Czech Science Foundation (No. 22-21903S) to V.B. and Italian Ministry of Health 5 × 1000 funds 2020 to P.M.

## Notes

### Competing Interest Statement

The authors have declared no competing interest.

