## Supplementary Materials for "The presence of a G-quadruplex prone sequence upstream of a minimal promoter increases transcriptional activity in the yeast *S. cerevisiae*"

### Supplementary Figures and legends

Libuše Kratochvilová[1,2,&], Matúš Vojsovič[1,2,&], Natália Valková[1], Lucie Šislerová[1,2], Zeinab El Rashed[3], Alberto Inga[4], Paola Monti[*5], Václav Brázda[1,2,*§]

§Corresponding author:

Václav Brázda

Institute of Biophysics of the Czech Academy of Sciences, Královopolská 135, 61265 Brno, Czech Republic

| Oligonucleotides | $I_{(0)}$ Tris-HCl | $I_{(KCl)}$ Tris-HCl + 100mM KCl | $I_{(KCl)}/I_{(0)}$ |
|---|---|---|---|
| **PUMA** | 1.75 ± 0.23 | 1.39 ± 0.04 | ↓ 0.79 |
| **KSHV** | 16.93 ± 0,08 | 19.22 ± 0.38 | ↑ 1.14 |
| **KSHV-1NO** | 25.36 ± 0.12 | 31.76 ± 0.44 | ↑ 1.25 |
| **KSHV-2NO** | 9.73 ± 0.61 | 25.69 ± 0.76 | ↑ 2.64 |
| **KSHV-Mut2.0** | 19.87 ± 0.87 | 23.89 ± 1.40 | ↑ 1.20 |
| **KSHV-Mut1.5** | 15.24 ± 1.20 | 10.14 ± 0.15 | ↓ 0,67 |
| **KSHV-3NO** | 6.98 ± 0.92 | 10.58 ± 0.73 | ↑ 1.51 |

**Table S1. Fluorescence intensity $I/I_0$ determined from oligonucleotides with the potential to form G4s .** The average fluorescence intensity of three repetitions was related to the fluorescence intensity of the blank (ThT with the appropriate buffer). The $I_{(KCl)}/I_{(0)}$ fold indicates the fold decrease (↓) or increase (↑) of fluorescence emission of samples in buffer with the addition of K+ ions compared to samples in Tris-HCl without KCl.

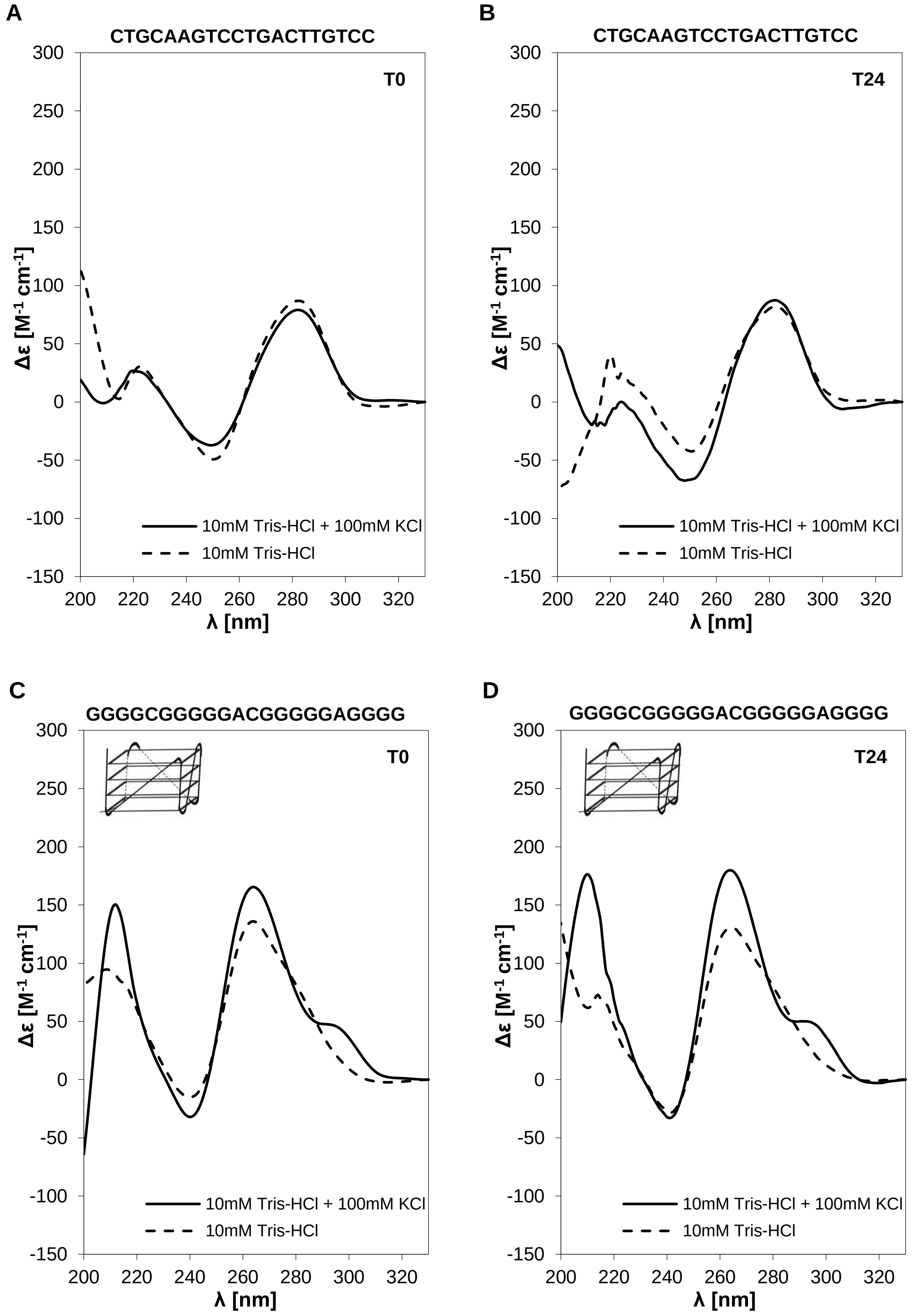
A
CTGCAAGTCCTGACTTGTCC
T0
Δε [M-1 cm-1]
λ [nm]
10mM Tris-HCl + 100mM KCl
10mM Tris-HCl
B
CTGCAAGTCCTGACTTGTCC
T24
Δε [M-1 cm-1]
λ [nm]
10mM Tris-HCl + 100mM KCl
10mM Tris-HCl
C
GGGGCGGGGGACGGGGGAGGGG
T0
Δε [M-1 cm-1]
λ [nm]
10mM Tris-HCl + 100mM KCl
10mM Tris-HCl
D
GGGGCGGGGGACGGGGGAGGGG
T24
Δε [M-1 cm-1]
λ [nm]
10mM Tris-HCl + 100mM KCl
10mM Tris-HCl

**E**

**GAAGCGGGGGACGGGGGAGGGG**

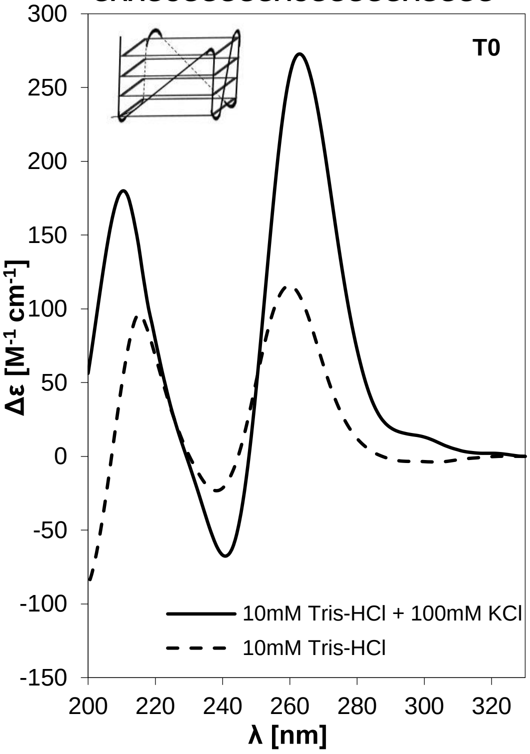


**F**

**GAAGCGGGGGACGGGGGAGGGG**

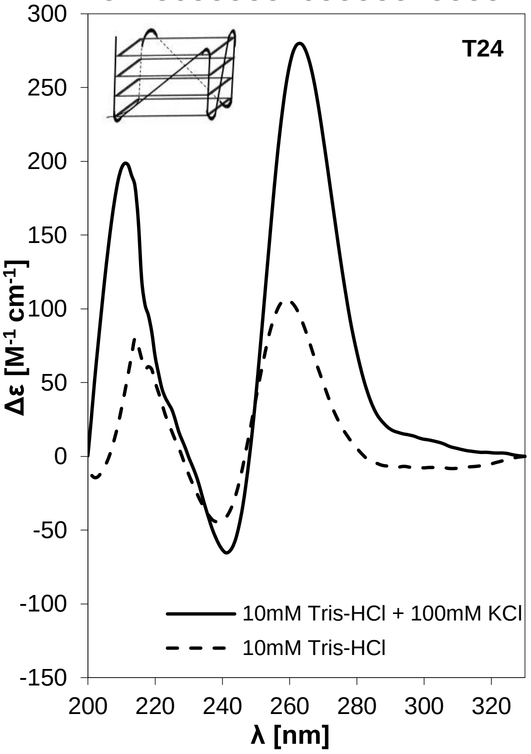


**G**

**GAAGCGAAAGACGGGGGAGGGG**

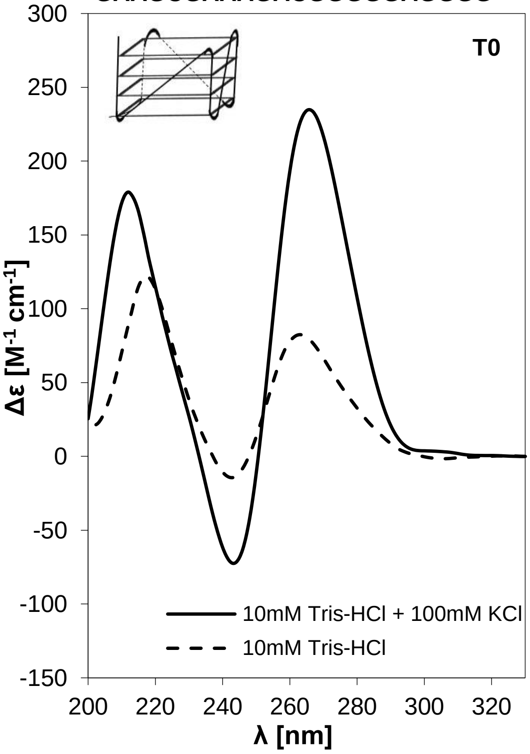


**H**

**GAAGCGAAAGACGGGGGAGGGG**

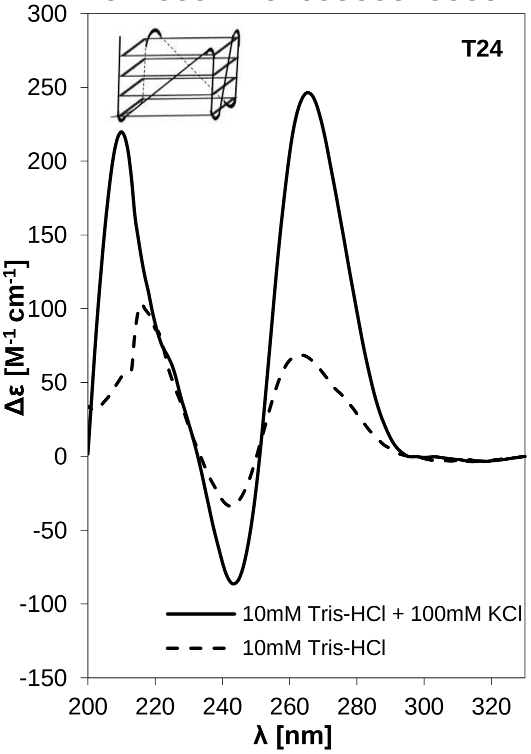

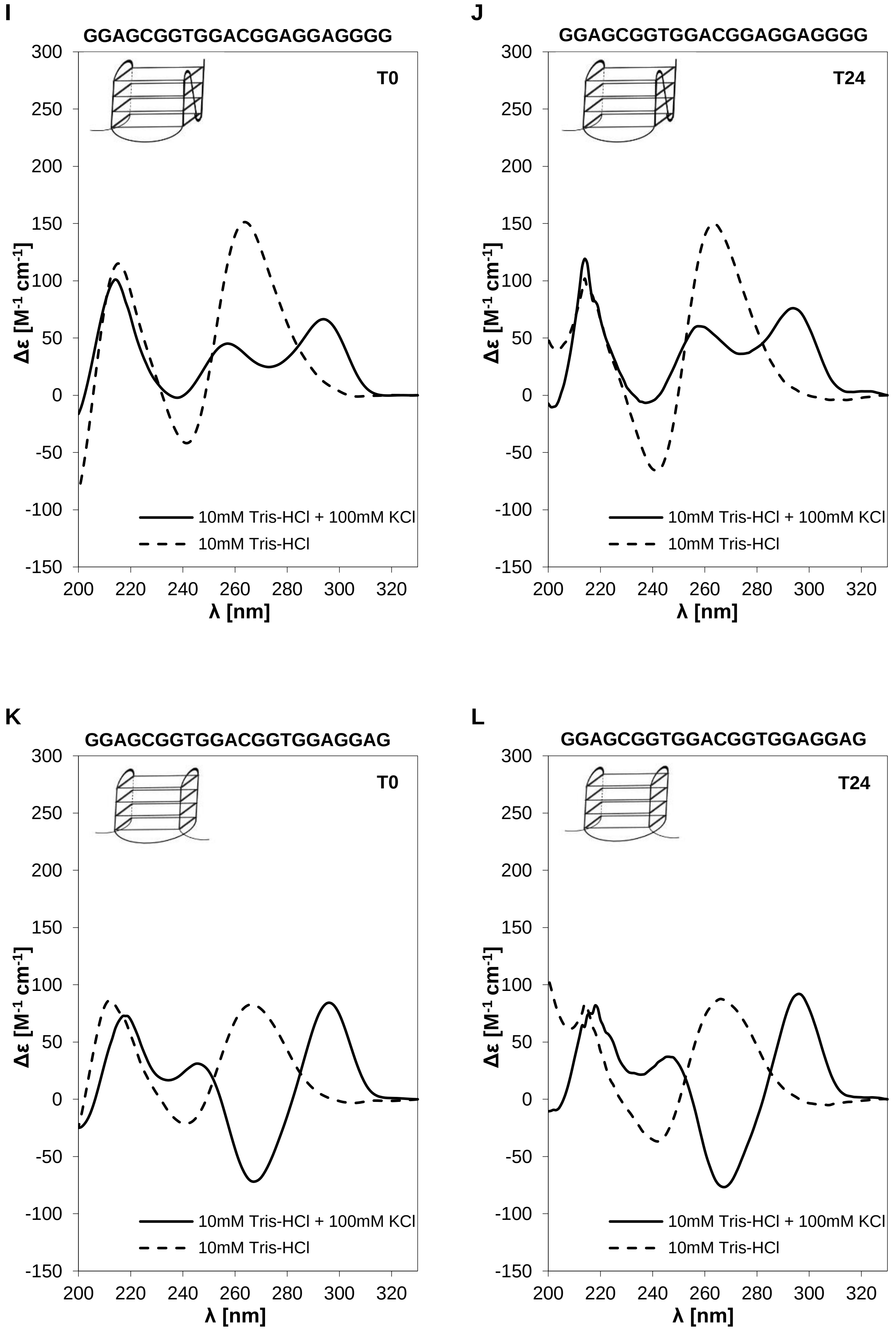

I
GGAGCGGTGGACGGAGGAGGGG
T0
J
GGAGCGGTGGACGGAGGAGGGG
T24
K
GGAGCGGTGGACGGTGGAGGAG
T0
L
GGAGCGGTGGACGGTGGAGGAG
T24
Δε [M-1 cm-1]
λ [nm]
10mM Tris-HCl + 100mM KCl
10mM Tris-HCl
300
250
200
150
100
50
0
-50
-100
-150
200
220
240
260
280
300
320

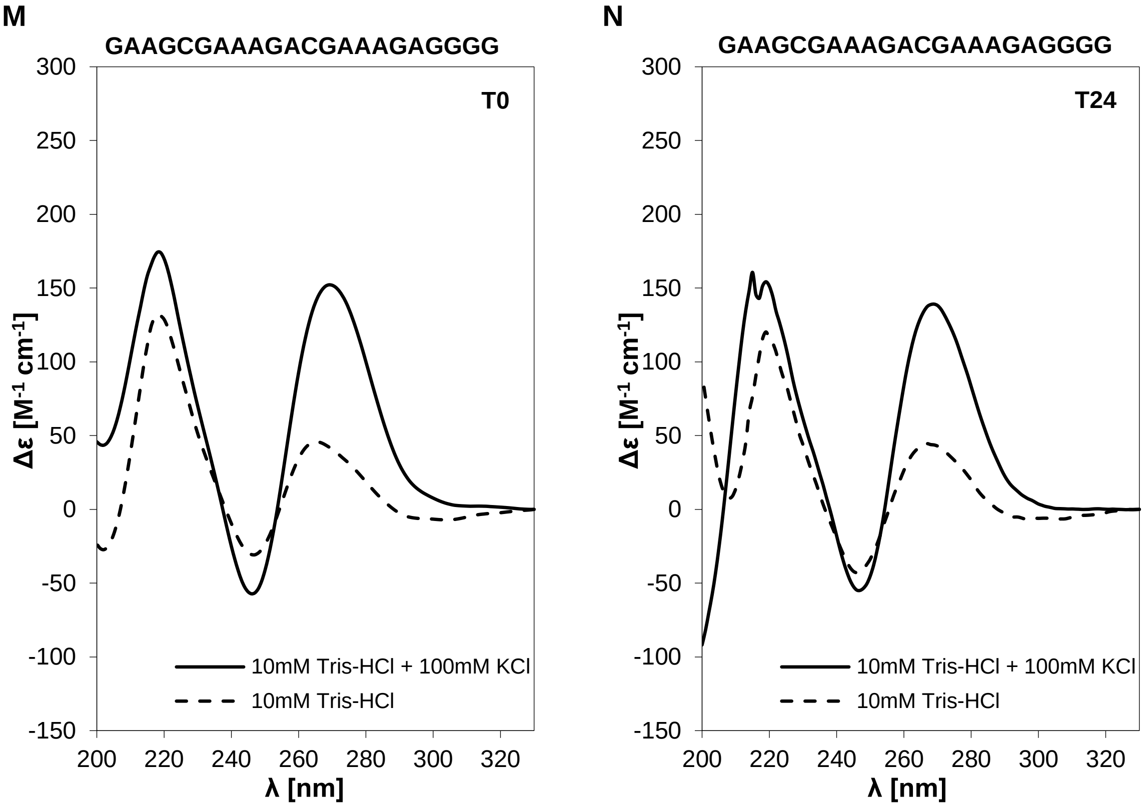


**Figure S1. CD spectra of oligonucleotides under study at $T_0$ (left panel) and $T_{24}$ (right panel).** CD spectra of PUMA oligonucleotide **(A, B)**, KSHV oligonucleotide **(C, D)**, KSHV-1NO oligonucleotide **(E, F)**, KSHV-2NO oligonucleotide **(G, H)**, KSHV-Mut2.0 oligonucleotide **(I, J)**, KSHV-Mut1.5 oligonucleotide **(K, L)** and KSHV-3NO oligonucleotide **(M, N)**. The solid line shows the spectra of the sample hybridized in 10 mM Tris-HCl with the addition of 100 mM KCl. The spectra of the sample hybridized in the medium without the addition of $K^+$ ions are plotted as a dashed line. The CD spectra at T0 from Figure 2 were also reproduced here to facilitate comparisons between the two time points.

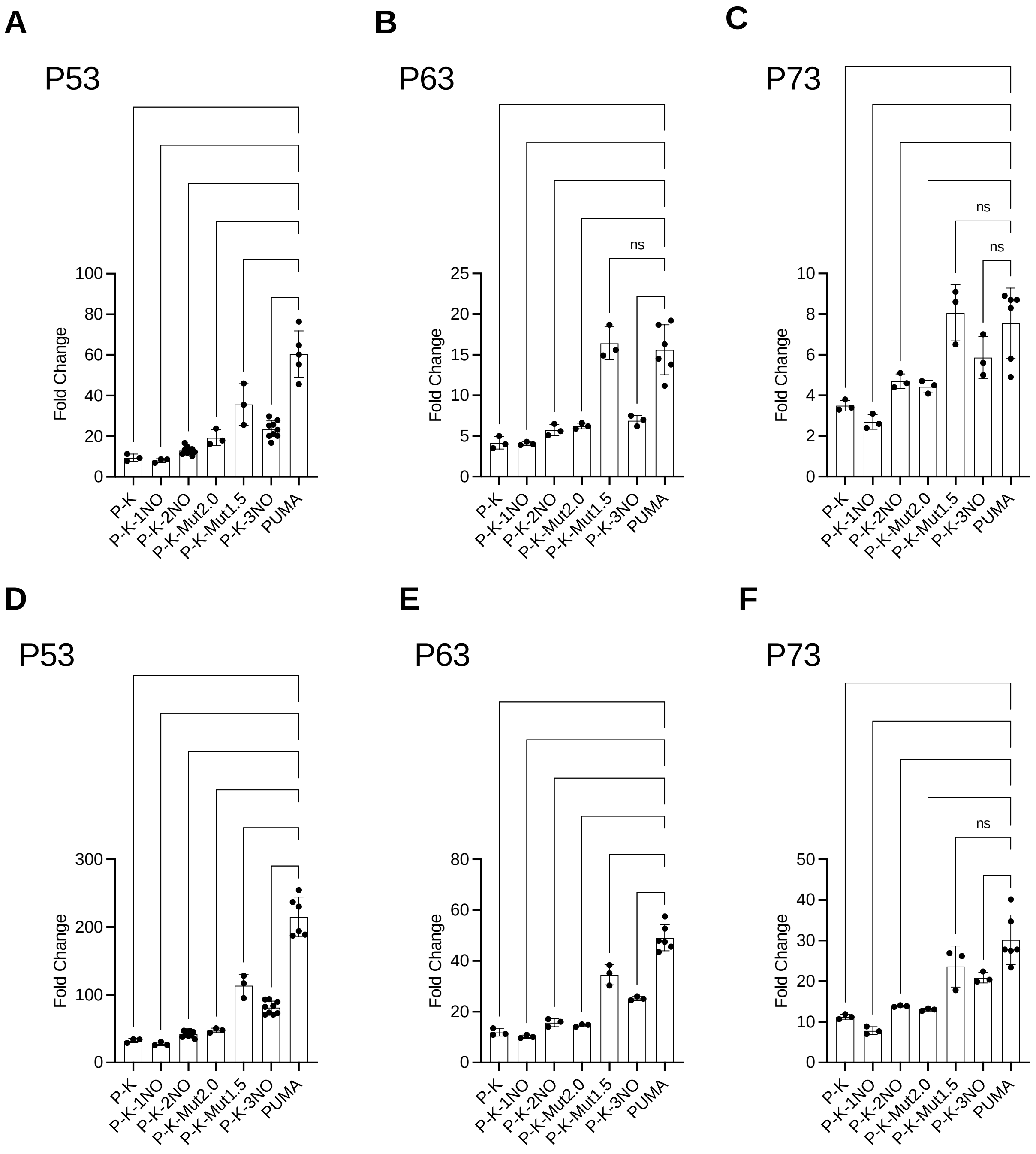


**Figure S2. Effect of G4 prone sequences on P53 family transactivation.** Fold change measurements for the indicated panel of yLFM reporter strains from P53, P63 and P73 yeast transformants at 0.016% galactose for 6 hours **(A-C)** or at 1% galactose for 6 hours (**D-F**)**.** Data are presented as mean ± standard deviation (SD) of at least three biological replicates. Individual values are also plotted. The symbols *, **, *** and **** indicate significant differences for p≤*0.0146*, p=0.0065, p=0.0006 and p<0.0001, respectively between PUMA strain and those containing other G4 regulatory elements. ns, not significant. Ordinary one-way ANOVA test.

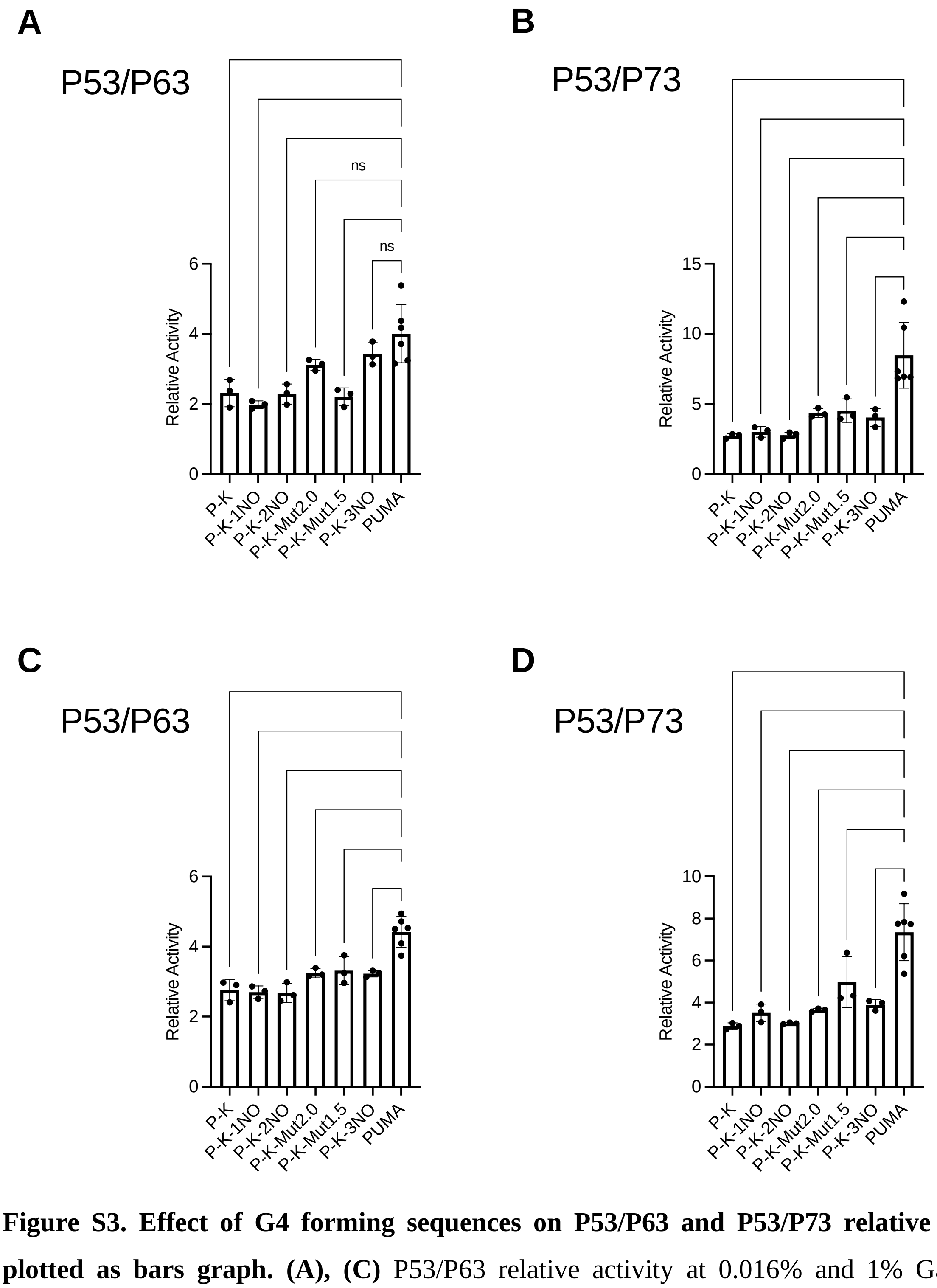


**Figure S3. Effect of G4 forming sequences on P53/P63 and P53/P73 relative activity plotted as bars graph. (A), (C)** P53/P63 relative activity at 0.016% and 1% Galactose, respectively in the indicated panel of yLFM reporter strains. **(B), (D)** P53/P73 relative activity at 0.016% and 1% Galactose, respectively as above. Data are presented as mean ± standard deviation (SD) of at least three biological replicates. Individual values are also plotted. The symbols **, *** and **** indicate significant differences for p*≤0.0064*, p=0.0009, and p<0.0001, respectively between PUMA strain and those containing other G4 regulatory elements. ns, not significant. Ordinary one-way ANOVA test.
